# Dynamic evolution of the major transcription factor DNA binding domain, and protein-protein interaction families during the evolution of the avian lineage

**DOI:** 10.1101/193896

**Authors:** Allie M. Graham, Jason S. Presnell

## Abstract

Transcription factors are characterized by their domain architecture, including DNA binding and protein-protein interaction domain combinations, which regulate their binding specificity, as well as their ability to effect a change on gene expression of their downstream targets. Transcription factors are central to organismal development, thus they potentially are instrumental in producing phenotypic diversity. Transcription factor abundance was estimated via 49 major DNA binding domain families, as well as 34 protein-protein interaction domain families, in 48 bird genomes, which were then compared with 6 available reptile genomes, in an effort to assess the degree to which these domains are potentially connected to increased phenotypic diversity in the avian lineage. We hypothesized that there would be increased abundance in multiple transcription factor domain families, as well as domains associated with protein-protein interactions, that would correlate with the increased phenotypic diversity found in birds; instead, this data shows a general loss/contraction of major domain families, with the largest losses in domain families associated with multiple developmental (feather, body-plan, immune) and metabolic processes. Ultimately, the results of this analyses represent a general characterization of domain family composition in birds, thus the specific domain composition of TF families should be probed further, especially those with the largest reductions seen in this study.

## INTRODUCTION

Transcription factors (TFs) are proteins that bind to DNA in a sequence-specific manner and enhance or repress gene expression. In response to a broad range of stimuli, transcription factors coordinate the regulation of gene expression of essential for defining morphology, functional capacity, and developmental fate at the cellular level. Although, transcription factor binding domains are very well conserved, the other associated domains, largely responsible for protein-protein interactions (PPI), readily diverge among homologs. Therefore, the structure and function of transcription factors are inherently modular. This attribute is thought to give gene-regulatory networks the ability to evolve more readily (Jarvela and Hinman 2015; Wray 2007), and could account for the seemingly large phenotypic difference between closely related groups (Liu et al. 2014; Nadimpalli et al. 2015).

A long-standing question has been whether changes in gene regulation or protein sequence have made a larger contribution to phenotypic diversity seen between species (Britten and Davidson 1969; King and Wilson 1975). Now, it is understood that changes in cis-regulatory systems more often underlie the evolution of morphological diversity than gene duplication/loss or protein function (Carroll 2008; Levine and Tjian 2003; Wittkopp and Kalay 2011). These cis-regulatory elements typically regulate gene transcription by functioning as binding sites for transcription factors. However, another avenue would be through whole-scale changes in transcription factor function via changes in domain modularity, either through their DNA-binding domains (DBD) and/or through other domains usually involved in protein-protein interactions (Schmitz et al. 2016; Wagner and Lynch 2008; Wagner and Lynch 2010), otherwise known as trans-regulatory elements. Phenotypic variation, from individual organisms to broad groups, has been attributed to a combination changes associated with cis- and trans-regulatory elements (Schmitz et al. 2016).

Although transcription factor diversity has been correlated with increased “complexity” across the eukaryotic lineage (Charoensawan et al. 2010; de Mendoza et al. 2013; Lehti-Shiu et al. 2016), no study has measured such transcription factor diversity within a specious, but highly related, animal clade. However, forty-eight avian genomes representing all the major families of birds (Aves) have recently been published (Eöry et al. 2015; OBrien et al. 2014; Zhang et al. 2014a), providing a unique opportunity to do just that. Birds represent one of the most diverse vertebrate lineages, and has the distinction of being the tetrapod class with the most living species, with half of them being Passerines (over 10,000 species) (Gill and Wright 2006). Not only do birds live worldwide and range in size (5 cm - 2.75 m), but they also variy widely in morphology, physiology and behavior, and have unique features (ie. feathers). From a genomic standpoint, these organisms have relatively low rates of gene gain/loss in gene families and have similarly sized genomes (0.91-1.3 Gb) (Zhang et al. 2014c).

Therefore, in this study, the major metazoan TF families were characterized by their major DNA-binding Domains (DBD), as well as protein-protein interaction (PPI) domains, in the 48 avian and 6 reptile genomes. This is the first study to analyze the evolutionary history and phylogenetic distribution of transcription factor domain families in the diverse genomes available for avian group, and the closely related reptile lineage. We hypothesized that there would be increased TF DBD or PPI abundance in multiple TF families, correlated with the increased phenotypic diversity found in birds; however, these results show wholescale losses within major TF DBD and PPI families, potentially reflecting general genome reductions seen between reptiles and birds. The largest losses in TF DBD families were associated with multiple developmental (feather, body-plan, immune) and metabolic processes.

## MATERIALS AND METHODS

### TF DBD Identification

Genomic data, in the form of protein models, were obtained complete genomes from publicly available databases for birds (previously at http://avian.genomics.cn/en/jsp/database.shtml; now at http://avian.genomics.cn/en/jsp/database.shtml) and reptiles (http://crocgenomes.org/). In general, these genomes were sequenced and assembled using similar techniques, and thus any systematic issues would be mitigated to an extent (see Zhang *et al*., 2014), though not eliminated. For example, half of the new avian genomes are considered low-coverage (24X – 39X), while the other half was considered high-coverage (50X – 115X)(Zhang et al. 2014b). In addition, three of the previously published genomes Chicken (*Gallus gallus*), Zebra finch (*Taeniopygia guttata*) and Turkey (*Meleagris gallopavo*) were sequenced using Sanger sequencing and are relatively low-coverage as well (7X – 17X).

A PfamScan was performed on the protein models (Finn et al. 2015), using a custom database containing only the (1) 49 major DBD families and (2) 34 protein-protein interaction domains. The “gathering threshold” option was selected in order to minimize false positives, though type of approach can underestimate total counts for some domains (Eddy 2011). The major DBDs and protein-protein interaction domains, were chosen based on previous studies (TD DBDs – de Mendoza et al., 2013), and if they additionally present in Chordates and available through PFAM/Interpro (PPI). For the major DBDs, in all cases, I defined a one-to-one relationship between TF class and DBD class (ie. Non-duplicates). DBDs that appeared just in combination with other DBDs (ie. Duplicates) were analyzed separately, to avoid an overestimation of TF numbers, due to problems detecting repeated domains (de Mendoza et al. 2013). For the PPI, no such designation was made as most PPI are combinatorial by nature.

### Assessment of TF DBD and PPI Gene family expansion/contraction

The enrichment of TF numbers was tested using a Mann-Whitney U test, with a significance threshold of *P* < 0.01, as similarly performed by de Mendoza et al. (2013). In addition, CAFE v4.0 (Computational Analysis of gene Family Evolution) was used to identify significantly expanding and contracting DBD families across Aves (De Bie et al. 2006). Because CAFE requires a ultrametric phylogenetic tree, the newick tree from Zhang et al (2014) was run through the program r8s (Sanderson 2003). CAFE was set to identify significantly expanding and contracting gene families, at a corrected significance level of p = 0.05, based on our data in conjunction with a Newick tree from Zhang et al. (2014a). The birth-death rate parameter, lambda, was determined with the -s flag that optimizes the log likelihood of data for all families. Several comparisons were made including (1) reptiles vs birds, (2) vocal-learning birds vs other birds (3) birds-of-prey vs other birds and (4) water-birds vs other birds.

### TF DBD and PPI Gene Ontology term characterization

To better understand the types of biological functions of the domains which were significantly different between birds and reptiles, gene ontology (GO) term analysis was performed using a representative member of birds with better GO annotations, chicken (*Gallus gallus*). Proteins with IDs containing the significant TF DBDs and the PPIs were extracted, and separately input into PANTHER to be compared against the rest of the genes currently annotated with the genome (FDR P < 0.05; PANTHER GO-Slim for Biological Process) (Mi et al. 2013a; Mi et al. 2013b). In addition, the same analysis was done with the domain families significantly different from the CAFE analyses.

## RESULTS

### Transcription Factor DBD Identification

In order to ascertain which families may have experienced family expansion or contraction, comparisons were made against the 6 reptile genomes (anole lizard, crocodile, alligator, gharial, green sea turtle, and soft-shelled turtle).

In the non-duplicate group, of the 49 DBD, 14 of them were significantly different (MWU, *P* > 0.01; Figure 1; Table 1; SUPP Table 1 and 2). Almost all of these families experienced a reduction in DBD presence, with an average decrease of 49.86%, while only 1 family showed an increase (Homeobox_KN), though the increase was substantial (844%; SUPP Table 2). However, it is important to note that most Homeobox_KN DBDs are associated with other DNA binding domains (“Duplicates”), which as a category, was not significantly different between reptiles and birds. MADF also showed a significant difference; however, this was driven by an extreme number in the soft-shelled turtle, and so was disregarded. In the duplicate group (ie. multiple DBD within a given protein), only 1 showed a significant difference – bZIP-MAF (Table 1). For the PPI domains, 79% (27 of the 34) showed a significant difference between reptiles and birds (Table 1).

**Figure 1:**
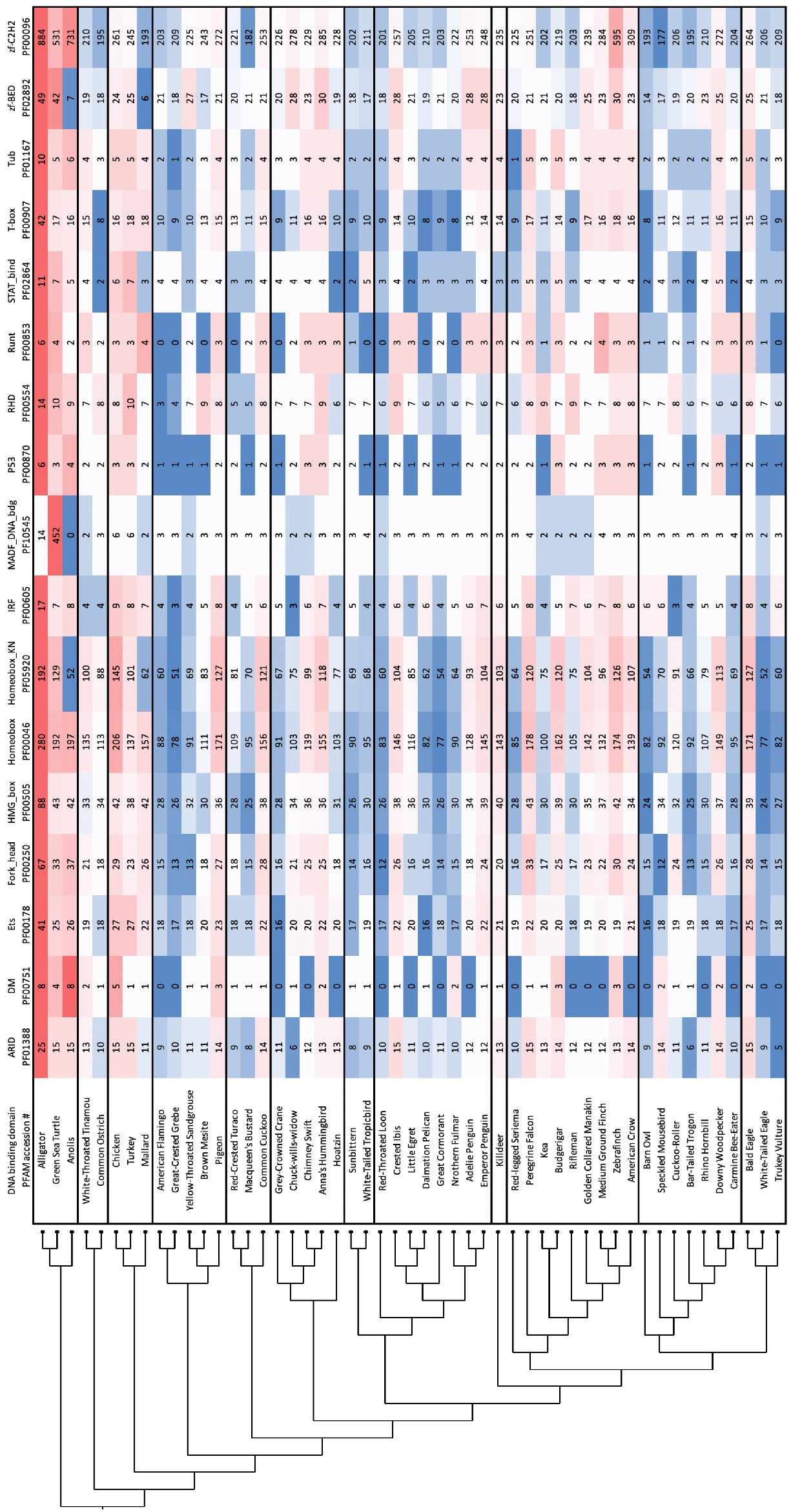
The 15 significantly different TF DBD families (duplicates, nonduplicates combined) against a phylogenetic tree of reptile representatives and the 48-bird species (red = more, blue = less)

**Table 1:** Non-duplicate DNA-Binding Domains (DBD) and Protein-Protein Interaction Domains (PPI) that are significantly different between reptiles, and birds (Mann-Whitney U test, *P*<0.01): “reptiles” represents the average number across the 6 lineages, and “birds” represents the average across the 48 lineages.

| Category | Domain Name | REPTILES | BIRDS | MWU | % Change |
| --- | --- | --- | --- | --- | --- |
| DBD | ARID | 15 | 10.75 | 0.00236 | -28.33% |
| DBD | DM | 4.333333333 | 0.8541667 | 0.00128 | -80.29% |
| DBD | Ets | 14.33333333 | 8.3541667 | 0.0005 | -41.72% |
| DBD | Fork_head | 37.33333333 | 19.6875 | 0.00016 | -47.27% |
| DBD | HMG_box | 49 | 32.708333 | 0.00046 | -33.25% |
| DBD | Homeobox | 75.16666667 | 32.583333 | 0.0002 | -56.65% |
| DBD | Homeobox_KN | 0.166666667 | 1.5625 | 0.00076 | 837.50% |
| DBD | IRF | 9.333333333 | 5.3958333 | 0.00104 | -42.19% |
| DBD | MADF_DNA_bdg | 101.3333333 | 2.9166667 | 0.00854 | -97.12% |
| DBD | P53 | 3.833333333 | 1.8333333 | 0.00164 | -52.17% |
| DBD | RHD | 9.5 | 6.9791667 | 0.00528 | -26.54% |
| DBD | STAT_bind | 6.166666667 | 3.5625 | 0.00148 | -42.23% |
| DBD | T-box | 19.83333333 | 11.5 | 0.0003 | -42.02% |
| DBD | Tub | 5.5 | 3.1666667 | 0.00496 | -42.42% |
| DBD | zf-BED | 8.833333333 | 2.875 | 0.00008 | -67.45% |
| DBD | zf-C2H2 | 521.1666667 | 209.9375 | 0.00016 | -59.72% |
| PPI | 4_1_CTD | 40.83333333 | 3.2916667 | 0.00034 | -91.94% |
| PPI | BAH | 23 | 6.6458333 | 0.00034 | -71.11% |
| PPI | BEN | 27 | 8.7708333 | 0.00308 | -67.52% |
| PPI | CARM1 | 2.333333333 | 0.4583333 | 0.0038 | -80.36% |
| PPI | Cdc6_C | 3.166666667 | 1.4375 | 0.0038 | -54.61% |
| PPI | CortBP2 | 7.666666667 | 4.0625 | 0.00128 | -47.01% |
| PPI | ESR1_C | 3.833333333 | 0.7291667 | 0.00052 | -80.98% |
| PPI | F-box | 111.6666667 | 37.916667 | 0.00008 | -66.04% |
| PPI | FACT-Spt16_Nlob | 14.33333333 | 4.4583333 | 0.0004 | -68.90% |
| PPI | FF | 0.666666667 | 0.0833333 | 0.0004 | -87.50% |
| PPI | Ge1_WD40 | 75.33333333 | 45.916667 | 0.00016 | -39.05% |
| PPI | IG | 630.8333333 | 233.5 | 0.00014 | -62.99% |
| PPI | KRAB | 162.6666667 | 4.8541667 | 0.0001 | -97.02% |
| PPI | L27_2 | 9.333333333 | 1.4791667 | 0.00124 | -84.15% |
| PPI | LRR_1 | 240 | 119.10417 | 0.00008 | -50.37% |
| PPI | LRR_4 | 424.5 | 177.0625 | 0.00008 | -58.29% |
| PPI | NHR2 | 13.66666667 | 3.25 | 0.0001 | -76.22% |
| PPI | OST-HTH | 8.5 | 2.6666667 | 0.0001 | -68.63% |
| PPI | PAH | 8.166666667 | 2.9375 | 0.00758 | -64.03% |
| PPI | PET | 9.833333333 | 4 | 0.00008 | -59.32% |
| PPI | PYRIN | 19.5 | 1.1041667 | 0.00018 | -94.34% |
| PPI | Serine_rich | 5.666666667 | 2.6458333 | 0.00056 | -53.31% |
| *PPI* | SIX1_SD | 8.833333333 | 2.875 | 0.0002 | -67.45% |
| *PPI* | SRCR | 36.66666667 | 20.083333 | 0.00188 | -45.23% |
| *PPI* | zf-CW | 11.5 | 2.7916667 | 0.00008 | -75.72% |
| *PPI* | zf-TFIIC | 3.166666667 | 1.0833333 | 0.00932 | -65.79% |

### DBD and PPI family expansion and contraction

CAFE was used to identify significantly expanding and contracting domain families (DBD, PPI), for several different comparisons within and between birds and reptiles, as well as within birds (water-bird, bird-of-prey, vocal-learners). Three of the 15 TF DBDs were unable to analyzed by CAFE due to memory issues caused by know program limitations, (ie. families having large numbers; Homeobox, Homeobox KN, zf-C2H2), so they were removed from analysis as per suggestion (CAFE Documentation, Mar 2017). Four of the DBD families showed a significant evidence of contraction (Forkhead, HMG box, T-box, zf-BED; Table 1; SUPP Table 1); three of those same families were also shown to be correlated with changes associated with being a bird-of-prey, water-bird and vocal-learner (Table 2). Ultimately, only 1 of the DBDs was unique to the reptile/bird comparison (T-box), and no unique DBDs were associated with any of the within-Aves comparisons.

**Table 2:**
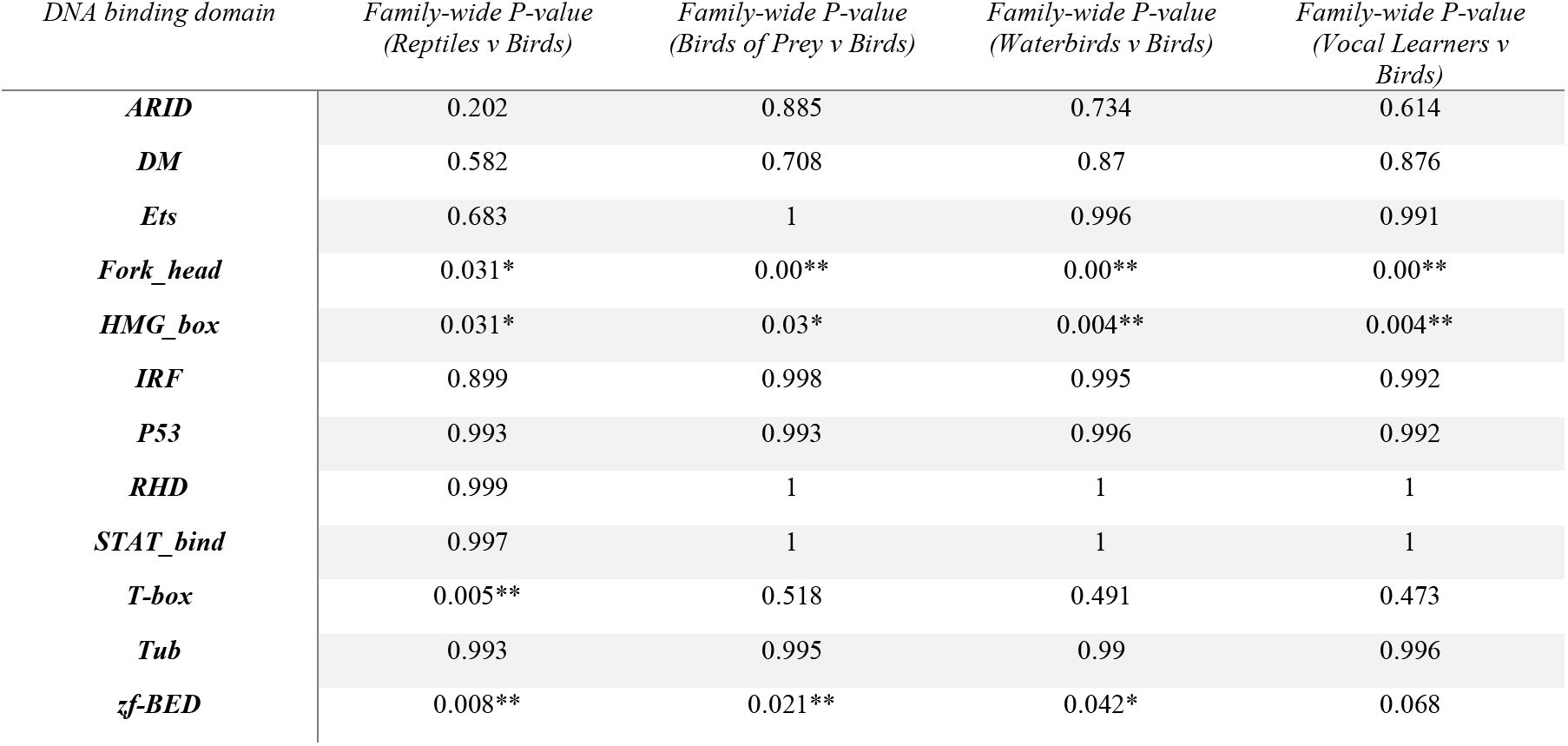
Expansion and contraction of TF DBD gene families in the avian lineages using the CAFE algorithm (asterixis indicate significance – * P-value < 0.05, ** P-value < 0.01)

Within the PPI domains, 5 were significant, 3 of those were unique to the reptile/aves compairison (BAH, BEN, SIX SD; SUPP Table 6, 7). One domain was significant across all comparisons (4.1 CTD), and one was significant for all within-Aves comparisons (SRCR; SUPP Table 2).

### Transcription Factor DBD and PPI Functional Characterization

For the comparison between reptiles and birds, the GO terms associated with the genes which contain members of the 14 TF DBD families are largely associated with metabolic processes (insulin receptor pathway, glucose homeostasis), and development (cell differentiation, embryonic development, skeletal development, anterior/posterior patterning) (Table 2; SUPP Table 4). These same families were identified by the CAFE analysis as having significantly contracted between reptiles and birds. Restricting the analysis to just those that were significant via the CAFE analysis, showed largely the same results (SUPP Table 5).

The terms related to the PPI domains were ones associated with more specific categories including histone deacetylation, collagen organization, actomyosin structure organization, as well as development (eye, sensory organ; SUPP Table 8, 9).

## DISCUSSION

The results of this analysis suggest there was an overall major reduction in both the TF DBD and PPI families across the avian lineage and showed few instances of substantial expansion. At present, although avian and reptile genomes are generally similarly sized, a reduction in genome size between reptiles and birds did occur in the saurischian dinosaur lineage between 230 and 250 million years ago (Organ et al. 2007). This coincided with a major reduction in repetitive elements, intron size, and even whole-scale loss of syntenic protein coding regions, typically attributed to the general metabolic requirements for flight (Lovell et al. 2014; Organ and Shedlock 2009; Wright et al. 2014; Zhang et al. 2014c; Zhang and Edwards 2012); thus, the reduction of TF DBD and PPI families seen in these results potentially mirrors the general genome reduction seen in other studies. However, it is somewhat surprising to not see any large instances of TF DBD family expansions, since increases in the number of regulatory proteins, including TFs, have frequently been connected to phenotypic innovations (Kusserow et al. 2005; Levine and Tjian 2003; Miyata and Suga 2001; Schmitz et al. 2016).

### Domain dynamics between reptiles and birds

It is interesting to note that the TF DBD families which experienced the sharpest declines were ones associated with heart development (T-box), vocal learning (Forkhead), feather formation (Ets), wing development, sex determination, and immune function (HMG box; Table 2,3); these same general results were also reflected in the most common GO terms represented (Table 2; SUPP Table 4). All of these aspects have been subjected to major physiological/developmental changes between reptiles and birds (Brusatte et al. 2015; Chatterjee 2015), especially the development of feathers from scales (Chuong et al. 2000), sex-determination through chromosomal differences rather than temperature (Sarre et al. 2004), and even changes in immune system functionality (Zimmerman et al. 2010). Although, it is difficult to speculate how reductions in these major families may be associated with such changes seen between avian and reptile lineages.

**Table 3:**
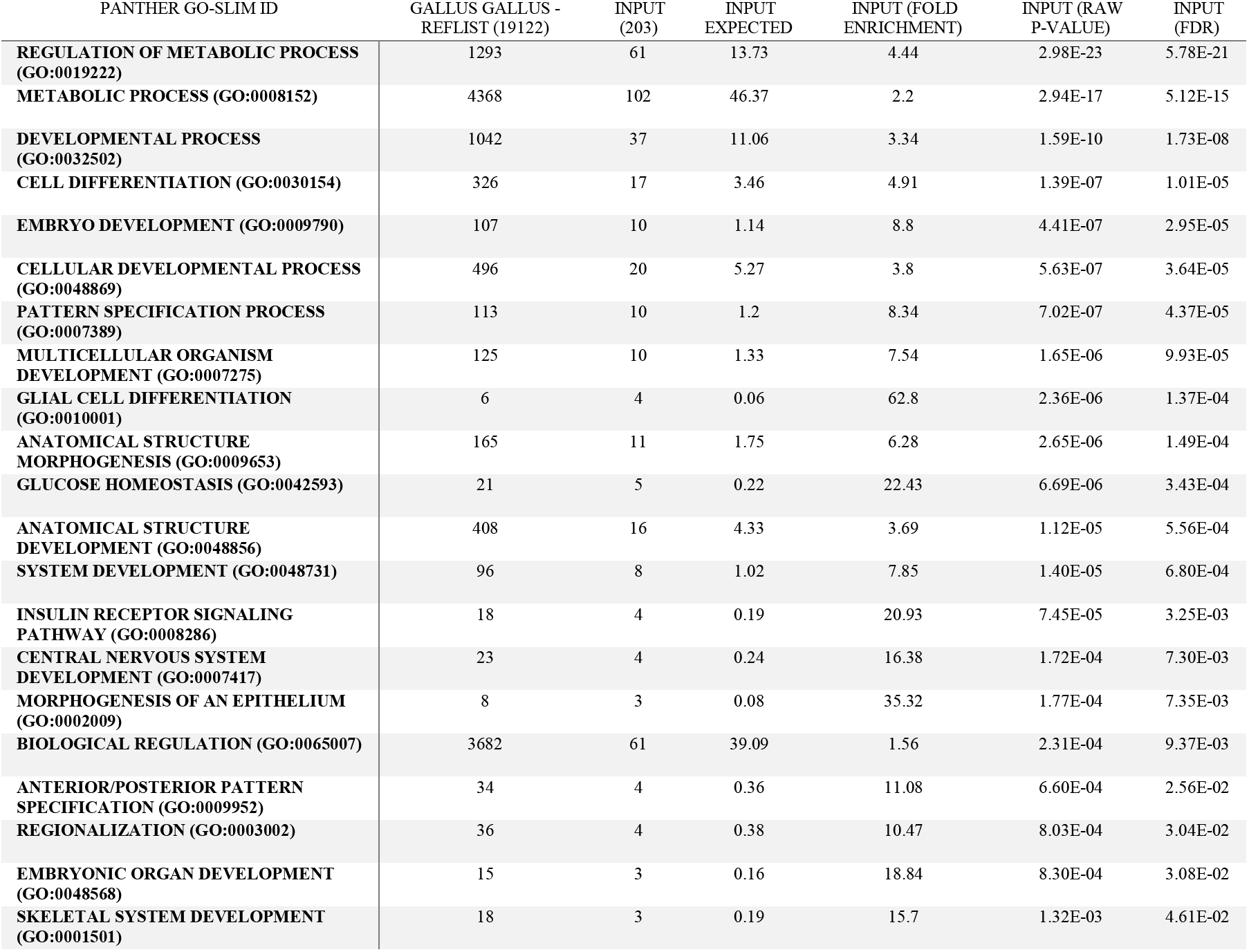
Gene Ontology analysis of Biological Processes using PANTHER for the 15 TF DBD families in Chicken (*Gallus gallus*): significant GO terms with FDR P < 0.05.

**Table 4:** Significantly different DBD categories, and their respective biological function (Mann-Whitney U; additional asterixis denote significance from CAFÉ analysis)

| *DOMAIN* | *DOMAIN "FUNCTION"* | *CITATION* |
| --- | --- | --- |
| *ARID* | general | PFAM: PF01388 |
| *DM* | sex determination/differentiation | (Smith et al. 2009)<br>InterPro: IPR001275 |
| *ETS* | feather formation | (Morgan et al. 1998) |
| *FORK_HEAD\** | general | (Hannenhalli & Kaestner 2009) |
|  | vocal learning | (Scharff & Haesler 2005) |
| *HOMEBOX* | general development | PFAM: PF00046 |
| *HOMEBOX_KN* | anterior-posterior axis formation | (Alonso 2002)<br>InterPro: IPR001356 |
| *HMG_BOX\** | wing development | (Welten et al. 2005) |
|  | sex determination | (Wallis et al. 2008) |
|  | immune function | (Lotze & Tracey 2005) |
| *IRF* | immune function | (Escalante et al. 1998)<br>SMART: SM00348 |
| *P53* | cell cycle (general) | InterPro: IPR011615 |
| *RHD* | immune/p53 | (Anrather et al. 2005)<br>InterPro: IPR011539 |
| *RUNT* | blood | (Kagoshima et al. 1993)<br>InterPro: IPR013524 |
| *STAT_BIND* | immune function | (Kisseleva et al. 2002)<br>InterPro: IPR012345 |
| *T-BOX\** | heart development | (Plageman & Yutzey 2005) |
| *TUB\** | cell cycle (general) | PFAM: PF01167 |
| *ZF-BED* | general | PFAM: PF02892 |
| *ZF-C2H2* | general | PFAM: PF00096 |

The PPI families which were significantly different between reptiles and birds were ones involved in apoptosis/inflammation (PYRIN), innate immune system (SRCR), chromatin structure (BAH), neural transcriptional repressors (BEN), development of sensory organs (SIX1 SD)(Dai et al. 2013; Fairbrother et al. 2001; Patrick et al. 2013; Sarrias et al. 2004; Yang and Xu 2013); the three latter of which were unique to the reptile-bird comparison, likely due to their typical co-occurrence with certain DBD families.

Therefore, the patterns seen in this study suggests that TF DBD and PPI families are largely not correlated with avian diversification; instead, this diversification could possibly be ascribed to other factors, such as protein-coding gene family duplications, or cis-regulatory changes (Seki et al. 2017; Zhang et al. 2014c). Duplications of protein-coding gene families are known to play a major role in species evolution: redundancy provides a medium for novelty while maintaining initial function (Lynch and Conery 2000; Zhang 2003). However, another possibility is that the absence of an increase TF DBD between reptiles and birds may instead suggest changes in TF modularity, appearing through increased interconnectivity/occurrence of DBD and PPI. Even though there was no increase in any particular PPI family, these domains could have potentially been re-organized/re-organized in conjunction with DBD families – such changes have been shown to alter their expression profiles or binding properties, in turn affecting the expression of many target genes, often with a major functional impact (Lespinet et al. 2002).

## CONCLUSTIONS

Ultimately, these results represent the first foray into TF DBD characterization between the avian and reptile genomes and suggests no link between TF DBD expansion and phenotypic diversity, in this case. Thus, these results of these analyses potentially strengthen the notion that cis-regulatory regions and protein-coding gene families are behind much of the extant avian diversification. Still, whole-scale reductions in TF DBD families in the genome likely posed a significant hurdle, unless these families were comprised of multiple members that were functionally redundant.

Overall, the results of this analyses represent a broad characterization of TF DBD family composition in birds; thus, the specific composition of TF families should be probed further, especially those with the largest reductions seen in this study. In addition, whether non-major TF DBD families have also seen a general reduction requires future analysis, as does the composition of domains associated with protein-protein interactions.

## Acknowledgements

We would like to thank the core facilities at Center for Genome Research and Biocomputing at Oregon State University, where certain elements of computing were performed. In addition, we would also like to thank the moderators at the CAFE Google Group, who were immensely helpful in troubleshooting the use of both CAFE and r8s.

## Data availability

Avian genomic data was pulled from its original repository location (http://gigadb.org/dataset/101000). The phylogenetic trees used in the CAFE analyses is available in its original repository location (http://gigadb.org/dataset/101041; WGT.ExaML.best.tre).

## Author contributions

AMG designed the study, AMG and JSP performed the data-collection, AMG analyzed the data and wrote the manuscript.

## Supplemental Information

<u>SUPP Table 1:</u> Full counts of DNA Binding Domain (DBD) families who are present only once in a given protein (ie. Non-Duplicates)

<u>SUPP Table 2:</u> Full counts of DNA Binding Domain (DBD) families who are present more than once in a given protein (ie. Duplicates)

<u>SUPP Table 3:</u> Full Gene Ontology (GO) terms for the genes from each DNA-Binding Domain (DBD) family (*Gallus gallus* reference)

<u>SUPP Table 4:</u> Overrepresented Gene Ontology (GO) terms for DNA-Binding Domains (DBD) which were significantly different between Reptiles and Birds (Mann-Whitney-U; *Gallus gallus* reference)

<u>SUPP Table 5:</u> Overrepresented Gene Ontology (GO) terms associated with DNA-Binding Domains (DBD) families that are significantly different between Reptiles and Birds via the CAFE analysis (*Gallus gallus* reference)

<u>SUPP Table 6:</u> Full counts for all protein-protein interaction domains (PPI) and their Mann-Whitney U test values (red = *P < 0.01*): “reptiles” represents the average number across the 6 lineages, and “birds” represent the average across the 48 lineages

<u>SUPP Table 7:</u> Results from the CAFE analysis for the protein-protein interaction domains (*P<0.05*): “reptiles” represents the average number across the 6 lineages, and “birds” represent the average across the 48 lineages.

<u>SUPP Table 8:</u> Overrepresented Gene Ontology (GO) terms for the genes containing the protein-protein interaction domains (PPI; *Gallus gallus* reference)

<u>SUPP Table 9:</u> Overrepresented Gene Ontology (GO) terms for the protein-protein interaction domains (PPI) that are significantly different between Reptiles and Birds via the CAFE analysis (*Gallus gallus* reference)

